# Calpain-mediated regulation of WAVE tyrosine dephosphorylation by PTP61F promotes lamellipodia formation

**DOI:** 10.64898/2026.10.08.757536

**Authors:** Wiebke Milani, Annalena Husmann, Sven Bogdan

## Abstract

WAVE is a major driver of lamellipodial protrusions. Previous studies suggested that Abelson (Abl)-dependent tyrosine phosphorylation of WAVE promotes actin polymerization and lamellipodia formation. Here, we provide evidence that Abl tyrosine kinase rather restricts lamellipodial localization of WAVE and inhibits lamellipodial cell spreading. Additionally, overexpression of a cytoplasmic PTP61F phosphatase increases WAVE localization at lamellipodia tips. PTP61F overexpression or pharmacological inhibition of Abl/Src kinases even protects cells from lamellipodia disruption induced by pervanadate, a potent protein tyrosine phosphatase inhibitor. By contrast, overexpression of a phosphatase-deficient PTP61F trapping mutant phenocopies loss of *wave* function. We identify the tyrosine Y234 as a prominent phosphorylation site in WAVE. Rescue experiments using CRISPR/Cas9-based KO wave mutant cells confirmed an important role of PTP61F-mediated WAVE dephosphorylation in lamellipodia formation. Our data suggest a model in which calpain-mediated proteolytic release of ER-bound PTP61F antagonizes Abl/Src-dependent WAVE phosphorylation and acts as a mechanism promoting re-localization of WAVE to lamellipodial tips.

## Introduction

The WAVE regulatory complex (WRC) is a key driver of Arp2/3-mediated actin filament branching and nucleation which is essential for forming sheet-like membrane protrusions known as lamellipodia in most eukaryotic cells (Chen et al., 2010). The WRC consists of five different protein subunits including Sra1/Cyfip1, Nap1/Hem2/Kette, Abi, HSPC300/BRICK1, and WAVE/SCAR (Rottner et al., 2021). The major activator of this hetero-pentameric complex is the small GTPase Rac which binds to the Sra1 subunit of the WRC and activates it by allosterically releasing the sequestered Arp2/3 activating VCA-domain of WAVE, allowing it to stimulate the Arp2/3-branched lamellipodial protrusions (Chen et al., 2017; Ding et al., 2022). A variety of additional upstream signals, including Arf GTPases, inositol phospholipids, membrane receptors, scaffolding proteins, and kinases might act cooperatively to relieve the WRC inhibition in the cell (Chen et al., 2014; Chen et al., 2010; Padrick et al., 2008). Both, WAVE and Abi proteins have many phosphorylation sites targeted by various tyrosine and serine kinases, such as Abl, Src, CK1/2 and Cdk5 (Krause and Gautreau, 2014; Mendoza, 2013). Serine phosphorylation of the VCA increases its affinity for the Arp2/3 complex and protects WAVE from degradation (Cory et al., 2003; Hirschhauser et al., 2021). By contrast, phosphorylation of the meander region of WAVE that binds the Sra1 subunit can directly destabilize VCA binding and evoke its activation (Chen et al., 2010). For example, *in vitro* studies showed that Abl-mediated phosphorylation of the mammalian WAVE complex in the meander region (Y151 in WAVE1/3, Y150 in WAVE2) relieved autoinhibition, exposing the VCA domain to activate the complex and induce lamellipodium formation (Chen et al., 2010; Leng et al., 2005; Stuart et al., 2006). Recent structure analysis further suggested a tyrosine lock mechanism involving Y151 in the meander sequence of WAVE1 that keeps the VCA domain of the WRC sequestered in the resting state (Ding et al., 2022). Binding of Rac1 promotes WRC activation by allosterically destabilizing this tyrosine lock region, which subsequently releases Y151 exposing it for phosphorylation by Abl. Once phosphorylated, Y151 cannot re-enter the tyrosine lock, even if Rac1 dissociates, thus effectively sustaining WRC activity (Ding et al., 2022). Thus, it appears that Rac1 and Abl tyrosine kinases act synergistically to drive WAVE-dependent actin polymerization. Paradoxically, several previous studies provided evidence that Abl tyrosine kinase rather inhibits cell migration and restrains lamellipodia extension during cell spreading (Jin and Wang, 2007; Kain and Klemke, 2001; Peacock et al., 2007). Moreover, inhibition of Abl stimulates tumor cell motility (Frasca et al., 2007; Marchal et al., 2023). In *Drosophila*, the protein tyrosine phosphatase PTP61F, the single orthologue of mammalian PTP1B and TCPTP acts as a direct antagonist of Abl in controlling lamellipodia formation (Huang et al., 2007; Ku et al., 2009). A model has been proposed in which a reciprocal regulation of Abi phosphorylation by Abl and PTP61F may regulate the localization and stability of the WRC (Huang et al., 2007). However, in this study ectopically expressed tagged Abi proteins were used that did not properly localized to lamellipodial tips like endogenous Abi but rather to punctate structures in cytoplasm (Huang et al., 2007). Abi is a known substrate of both Abl and PTP61F (Huang et al., 2007; Stephan et al., 2008), the physiological role of its tyrosine phosphorylation in lamellipodia formation, however, remains unknown.

Tyrosine phosphoproteomics using a cytoplasmic PTP61F (D203A) trapping mutant protein identified not only Abi but also other WRC components including Sra-1, Kette and WAVE itself as potential PTP substrates in cultured *Drosophila* S2 cells (Chang et al., 2008). However, a regulation of WAVE by Abl and PTP61F has not further been investigated. Like its mammalian orthologue, PTP61F localizes to the cytoplasmic face of the endoplasmic reticulum and acts as a negative regulator of several highly conserved signaling pathways, including the JAK–STAT pathway, the Insulin-like Receptor (INR) pathway the EGFR-pathway, and the platelet-derived growth factor (PDGF)- and vascular endothelial growth factor (VEGF)-receptor-related (PVR) pathway (Liu et al., 2022; Read and Wilson, 2024). In human platelets, cleavage of PTP1B C-terminus by the calcium-dependent protease calpain *in vivo* leads to the generation of a cytoplasmic, active enzyme that correlates with the transition from reversible to irreversible platelet aggregation (Frangioni et al., 1993). Cleavage and subcellular relocation of PTP1B/PTP61F might represent a common mechanism for altering phosphorylation of several known PTP1B substrates with important physiological implications in cell types other than platelets as previously shown for a calpain/PTP1B negative feedback regulation of VEGFR2 in endothelial cells (Zhang et al., 2017).

Here, we used cultured *Drosophila* S2R+ cells as an efficient model system to analyze the role of Abl and PTP61F in regulating WAVE localization and lamellipodia formation in more detail. Our data confirm that both Abl and PTP61F physically interact with WAVE. Different from previous studies, we found that genetic and pharmacological inhibition of Abl rather promotes lamellipodial cell spreading by the stabilization of WAVE at the leading edge whereas overexpression of a phosphatase-deficient PTP61F-D203A trapping mutant phenocopies loss of *wave* function. Correspondingly, overexpression of a truncated cytoplasmic PTP61F-DC variant lacking the ER localization motif and the coiled-coil region increases WAVE localization at lamellipodia tips promoting cell spreading. Rescue experiments with distinct phospho-mutant WAVE variants further suggest that the balance between Abl-mediated phosphorylation and PTP61F-mediated dephosphorylation at Y234 controls WAVE localization and thereby WAVE-dependent lamellipodia formation. In summary, we suggest a pivotal role for calpain in proteolytic cleavage of ER-bound PTP61F that promotes WAVE dephosphorylation and increased translocation to the cell edge.

## Results

### Suppression of *Drosophila* Abl function promotes lamellipodial WAVE localization and cell spreading

Previous studies in mammalian cell culture and in flies revealed that the non-receptor tyrosine kinase Abl interacts with and phosphorylates WAVE proteins stimulated by its positive interactor Abi, which is also a central WRC subunit (Juang and Hoffmann, 1999; Leng et al., 2005; Stephan et al., 2008; Westphal et al., 2000). We confirmed that *Drosophila* WAVE becomes highly tyrosine-phosphorylated upon treatment with the tyrosine phosphatase inhibitor pervanadate (Figure 1A). In western blots, an increase in tyrosine phosphorylation of endogenous WAVE upon pervanadate treatment was evident as a strong shift in electrophoretic mobility (Figure 1A). A similar shift in electrophoretic mobility was observed upon overexpression of constitutively active BCR-Abl kinase (Figure 1B, C; Zhuang et al., 2011). Immunoprecipitation experiments further confirmed that endogenous WAVE protein becomes highly-tyrosine phosphorylated upon overexpression of BCR-Abl kinase (Figure 1C).

**Figure 1.**
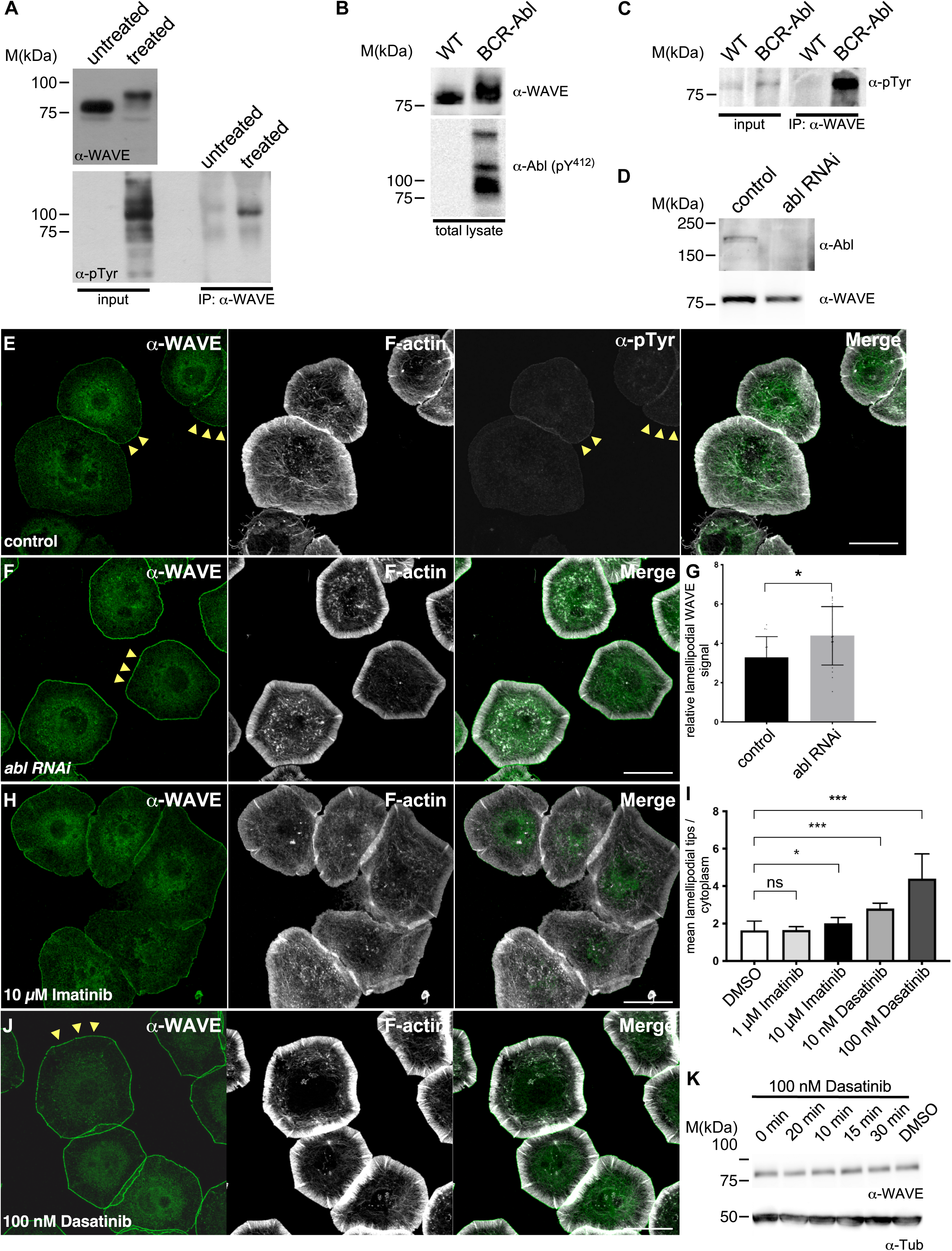
Inhibition of Abl and Src kinases increase WAVE localization at lamellipodial tips. **(A)** Western blot analysis of cell lysates from S2R+ cells untreated or treated with 1mM pervanadate resulting in a strong electrophoretic mobility shift of endogenous WAVE. Immunoprecipitation of WAVE confirmed pervanadate-induced hypertyrosine phosphorylation of WAVE, probed with anti-pTyr and anti-WAVE antibodies. **(B)** Expression of activated Abl (BCR-Abl) also results into a prominent mobility shift of endogenous WAVE. Activated BCR-Abl is detected by an antibody that recognizes the phosphorylated Y412-Abl (α-P^Y412^). **(C)** Immunoprecipitated endogenous WAVE is highly phosphorylated by BCR-Abl. **(D)** Western blots of lysates from S2 cells depleted for *abl*, probed with an anti-Abl or anti-WAVE confirmed a reduction of Abl expression. No significant changes in WAVE expression were observed. **(E)** Confocal images of S2R+ cells plated on concanavalin A, stained for endogenous WAVE (green), phosphotyrosine (pTyr, white) antibody and F-actin (Phalloidin, white). Scale bars = 20 µm. Arrowheads mark WAVE and phosphotyrosine immunostaining enriched at the leading edge. **(F)** RNAi-meditated depletion of *abl* leads to an increase of endogenous WAVE at lamellipodial tips (arrowheads). **(G)** Quantification of lamellipodial WAVE signal in *abl*-depleted cells compared to untreated control cells. Results are the average of three independent experiments. *P*-values from unpaired *t*-test are shown when significantly different from control (\**P*<0.05). Error bars represent standard deviation (SD). Control cells n=20; *abl* RNAi n= 19 **(H)** An increase of lamellipodial WAVE signal was also observed after treatment with 10 µM imatinib which specifically inhibits the Abl tyrosine kinase. **(I)** Quantification of increased lamellipodial/cytoplasm WAVE signal ratios in cells treated with imatinib and dasatinib at different concentrations. **(J)** A more dramatic increase of lamellipodial WAVE signal was observed after treatment with 100 nM dasatinib, a dual Src/Abl tyrosine kinase inhibitor. **(K)** Time-course western blot analysis shows no significant effect on overall endogenous WAVE protein level after treatment with 100 nM dasatinib. Tubulin served as loading control.

To analyze the function of Abl in WAVE-dependent lamellipodial cell spreading we knocked down Abl expression by RNAi in *Drosophila* S2R+ cells (Figure 1D), an efficient model for WAVE-dependent lamellipodia formation (Bogdan and Klämbt, 2003; Kunda et al., 2003; Rogers et al., 2003). Control S2R+ cells showed broad circumferential lamellipodia when spread on Concanavalin A (ConA) (Figure 1E; Bogdan et al., 2005; Kunda et al., 2003). Endogenous WAVE protein was enriched at the tips of lamellipodia in front of lamellipodial actin filaments, but a significant pool of WAVE protein was also found in the cytoplasm (Figure 1E, yellow arrowheads) (Bogdan et al., 2005; Kunda et al., 2003). RNAi-mediated knockdown of Abl did not disrupt lamellipodia formation (Figure1D, F). By contrast, cells depleted for *abl* showed an enhanced cell spreading and increased WAVE localization at lamellipodial tips (Figure 1F, yellow arrowheads). Quantification of immunostainings confirmed a significant increase of lamellipodial WAVE signal intensity up to 30 % at the expense of cytoplasmic localization compared to control cells (Figure 1G). Similarly, pharmacological treatment of S2R+ cells with the Abl inhibitor imatinib/STI-571 (Schindler et al., 2000), which has a high affinity for Abl kinase also led to a significant enrichment of the endogenous WAVE protein at lamellipodia tips (Figure 1H). Spreading experiments with S2R+ cells in the presence of the dual Abl/Src inhibitor dasatinib (Shah et al., 2004) in a nanomolar range resulted in an even more dramatic accumulation of the endogenous WAVE protein at lamellipodia tips at the expense of its cytoplasmic pool (Figure 1I, J). Western blot analysis of cell lysates from treated cells showed no changes of total WAVE protein (Figure 1K).

Tyrosine phosphorylation by Abl and Src might have synergistic effects on WAVE localization or activity. Overexpression of wild type GFP-tagged Abl did not affect S2R+ cell morphology. Abl-GFP is enriched at the leading edge of the cell. Tyrosine phosphorylation is increased in cells overexpressing Abl-GFP, whereas localization of endogenous WAVE is slightly reduced at lamellipodial tips. By contrast, overexpression of both constitutively active Abl (BCR-Abl) and constitutively active Src42A-CA disrupted proper lamellipodia formation (Figure 2B, C). Transfected cells showed induction of long filopodia-like protrusions, particularly upon overexpression of constitutively active Src42 kinase (Figure 2B, C). Taken together, these data strongly suggest that tyrosine phosphorylation is not essential for the initiation or formation of lamellipodia, but rather functions to modulate or optimize cellular protrusion efficiency.

**Figure 2.**
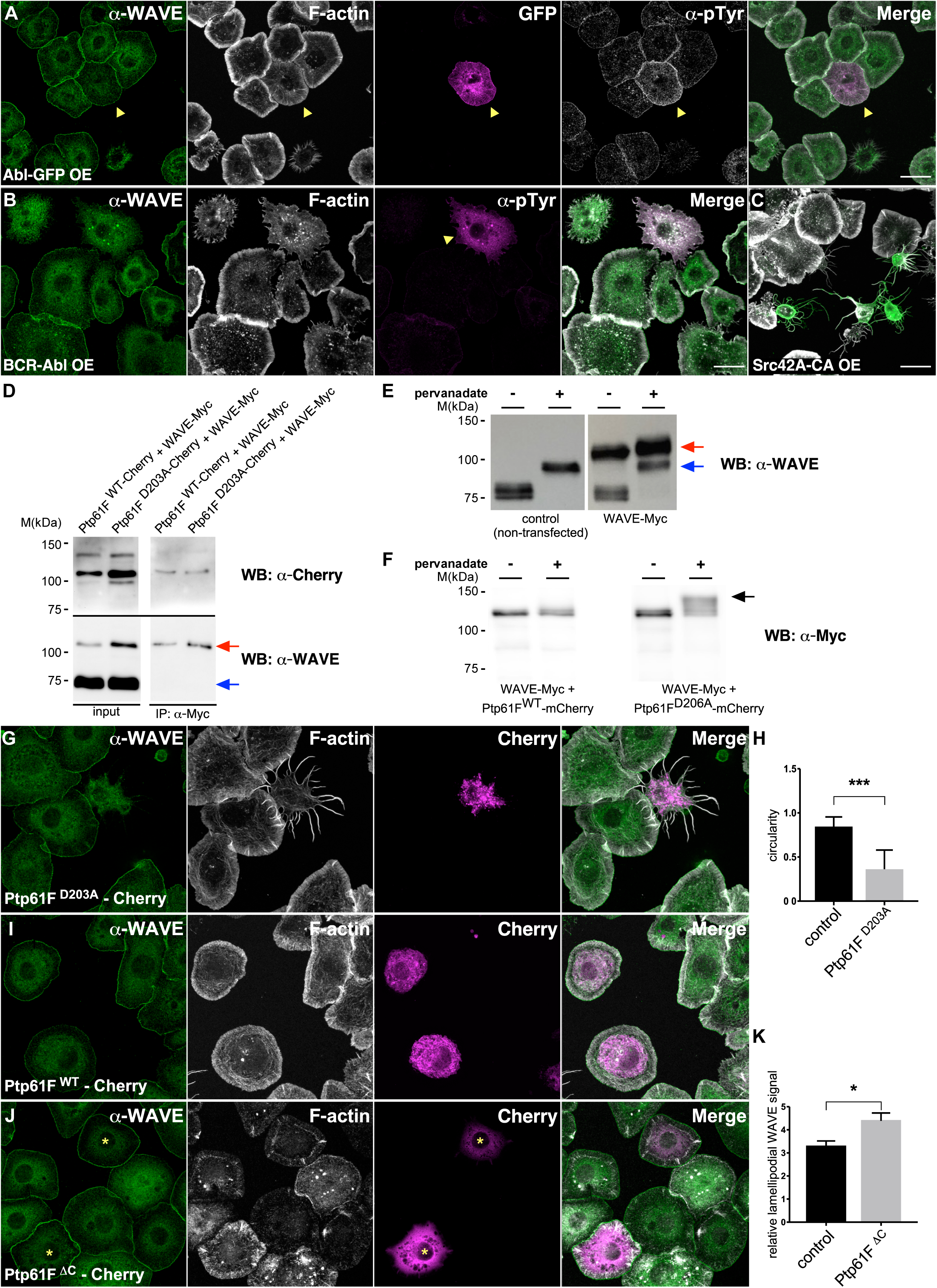
PTP61F interacts with WAVE and regulates WAVE-dependent lamellipodia formation. **(A)** Expression of GFP-tagged wild type Abl did not affect S2R+ cell morphology. Abl-GFP is enriched at the leading edge of the cell. Tyrosine phosphorylation is increased in cells overexpressing Abl-GFP, whereas localization of endogenous WAVE is slightly reduced at lamellipodial tips. **(B)** Overexpression of activated BCR-Abl dramatically induces tyrosine phosphorylation that disrupts lamellipodia formation. **(C)** Overexpression of activated Src-42A completely disrupts S2R+ cell morphology and induces long thin cellular protrusions. **(D)** Co-immunoprecipitation experiments with extracts of *Drosophila* S2R+ cells co-transfected with HA-tagged WAVE and either with a full-length Myc-tagged PTP61Fm or with a Myc-tagged substrate trapping mutant PTP61FmD203A that stably associates with a phosphorylated substrate. Full-length PTP61Fm and the PTP61FmD203A trapping mutant bound weakly to WAVE. The trapping mutant PTP61Fm-D203A binds more strongly WAVE than wild type PTP61Fm-WT protein. Overall, the interaction between PTP61F and WAVE was not further enhanced by treating the cells with 1mM pervanadate **(E)** Overexpression of PTP61F^D203A^ tagged with fluorescent Cherry tag abrogated lamellipodial cell spreading and induced a spiky morphology, similar to that observed in *wave* loss-of-function. **(F)** Quantification confirmed that PTP61Fm^D203A^ overexpression strongly reduced cell circularity. Results are the average of three independent experiments. Asterisks indicate significant P-values (*** < 0.001). Error bars represent standard deviation (SD). PTP61Fm WT n= 31; PTP61Fm^D203A^ n=12. **(G)** Overexpression of full-length PTP61Fm tagged with fluorescent Cherry tag did not induce filopodial-like protrusions. **(H)** Overexpression cytoplasmic PTP61FmDC protein tagged with fluorescent Cherry tag resulted in a prominent increase of endogenous WAVE at lamellipodia tips. Transfected cells are marked by an asterisk. **(I)** Quantification confirmed a prominent increase of endogenous WAVE at lamellipodia tips in cells expressing cytoplasmic PTP61F-ΔC compared to non-transfected control cells. Results are the average of three independent experiments. Asterisks indicate significant P-values (*** < 0.001). Error bars represent standard deviation (SD). Control cells n=21; PTP61F-ΔC n=27.

### The tyrosine phosphatase PTP61F interacts with WAVE and reduces its phosphorylation

Previous data suggest that the balance of WAVE tyrosine phosphorylation is a regulatory mechanism in actin-driven lamellipodia protrusions which might be tightly controlled by the reciprocal actions of Abl kinase and antagonistic protein tyrosine phosphatases (PTPs). Tyrosine phospho-proteomics already identified WAVE as a possible substrate of the protein tyrosine phosphatase PTP61F, the single orthologue of mammalian PTP1B and TCPTP and known antagonist of Abl functions in *Drosophila* (Chang et al., 2008; Huang et al., 2007; Ku et al., 2009). Alternative splicing of the PTP61F can result in two variants that only differ at their extreme C-termini. The splice variant PTP61Fm has a hydrophobic C-terminal tail similar to PTP1B, and is targeted to intracellular membranes including the ER, whereas the PTP61Fn variant lacks the hydrophobic C-terminus, and is targeted to the nucleus by a bipartite nuclear localization signal (NLS) like its mammalian TCPTP counterpart (Buszard et al., 2013; McLaughlin and Dixon, 1993).

We previously performed a yeast two-hybrid approach using the N-terminal WHD and the basic region of WAVE as a bait (Höhne et al., 2025), and also identified one clone encoding the ER-bound PTP61Fm isoform. To validate an interaction between PTP61Fm and WAVE we performed co-immunoprecipitation experiments with extracts of *Drosophila* S2R+ cells co-transfected with HA-tagged WAVE and either with a full-length Myc-tagged PTP61Fm-WT or with a Myc-tagged substrate trapping mutant PTP61Fm-D203A that stably associates with a phosphorylated substrate but does not dephosphorylate it (Figure 2D, F; Chang et al., 2008). We found that the trapping mutant PTP61Fm-^D203A^ interacts more strongly with WAVE than wild type PTP61Fm-^WT^ protein (Figure 2D). Overall, the interaction between PTP61F and WAVE was not further enhanced by treating the cells with pervanadate Figure 2D).

### Overexpression of phosphatase-dead PTP61Fm-D203A trapping mutant phenocopies loss of *wave* function

We further analyzed the function of PTP61F in WAVE-dependent lamellipodial cell spreading. RNAi-meditated knock down of PTP61F showed no defects, neither in S2R+ cell morphology nor in WAVE localization (Supplementary figure S1A, B). RNAi specificity was confirmed by using a transfected PTP61F-mCherry fusion which was efficiently downregulated in contrast to control proteins like Tubulin (Supplementary figure S1C).

We next analyzed the cellular consequences of different PTP61F constructs on lamellipodia formation. Interestingly, overexpression of PTP61F-D203A abrogated lamellipodial cell spreading and induced a spiky morphology (Figure 2E). Quantitative analysis confirmed that PTP61Fm-D203A overexpression strongly reduced cell circularity, thus phenocopied loss of WAVE function (Bogdan and Klämbt, 2003; Figure 2F). By contrast, expression of full-length PTP61Fm did not induce filopodial-like protrusions (Figure 2G). As previously reported (Buszard et al., 2013), PTP61Fm tagged with fluorescent Cherry tag localized predominantly at the perinuclear endoplasmic reticulum (Figure 2G). Removal of the hydrophobic C-terminal tail including the ER localization motif resulted in a complete cytoplasmic localization of PTP61Fm (PTP61Fm-ΔC, Figure 2H, yellow asterisks). Remarkably, expression of this cytoplasmic PTP61Fm-ΔC protein resulted in a significant increase of endogenous WAVE at lamellipodia tips (white asterisks Figure 2H, quantification in Figure 2I). Thus, overexpression of this cytoplasmic PTP61Fm-ΔC phenocopies loss of *abl* function.

### Expression of PTP61FmΔC protects cells from pervanadate-induced lamellipodia disruption

The cell-permeable reagent pervanadate is widely used to study phosphotyrosine signaling because it is a potent inhibitor of protein tyrosine phosphatases (Bennett et al., 1993). A more recent study showed that pervanadate also modulates phosphotyrosine signaling by oxidizing specific cysteine residues on Src tyrosine kinases, thereby leading to their hyperactivation (Mulholland et al., 2026). Both effects could contribute to rapid actin-dependent morphological changes following pervanadate treatment (Bennett et al., 1993). *Drosophila* S2R+ cells that fully spread on ConA were treated with 1mM pervanadate also showed prominent morphological changes (Figure 3A). Treatment with pervanadate typically disrupted lamellipodial protrusions within the first 20 minutes, accompanied by a re-localization of endogenous WAVE from the leading edge to cytoplasm. This effect was significantly suppressed by co-treatment with Abl/Src kinase inhibitor dasatinib suggesting that pervanadate-induced re-localization of endogenous WAVE might partially depend on Src hyperactivation (Figure 3C, D). Remarkably, pervanadate-induced lamellipodia disruption and WAVE re-localization was also completely suppressed in cells expressing the cytoplasmic PTP61Fm-ΔC variant (Figure 3E, F). Thus, binding of PTP61Fm-ΔC to WAVE appeared to protect the cells from the effects caused by pervanadate. This suggest that binding of PTP61Fm-ΔC to WAVE might restrict tyrosine phosphorylation of WAVE and subsequently protects from pervanadate-induced hyperphosphorylation. Supporting this notion, we found that cells expressing PTP61Fm-ΔC show a significant decrease in phosphotyrosine immunostaining (Figure 3G)

**Figure 3.**
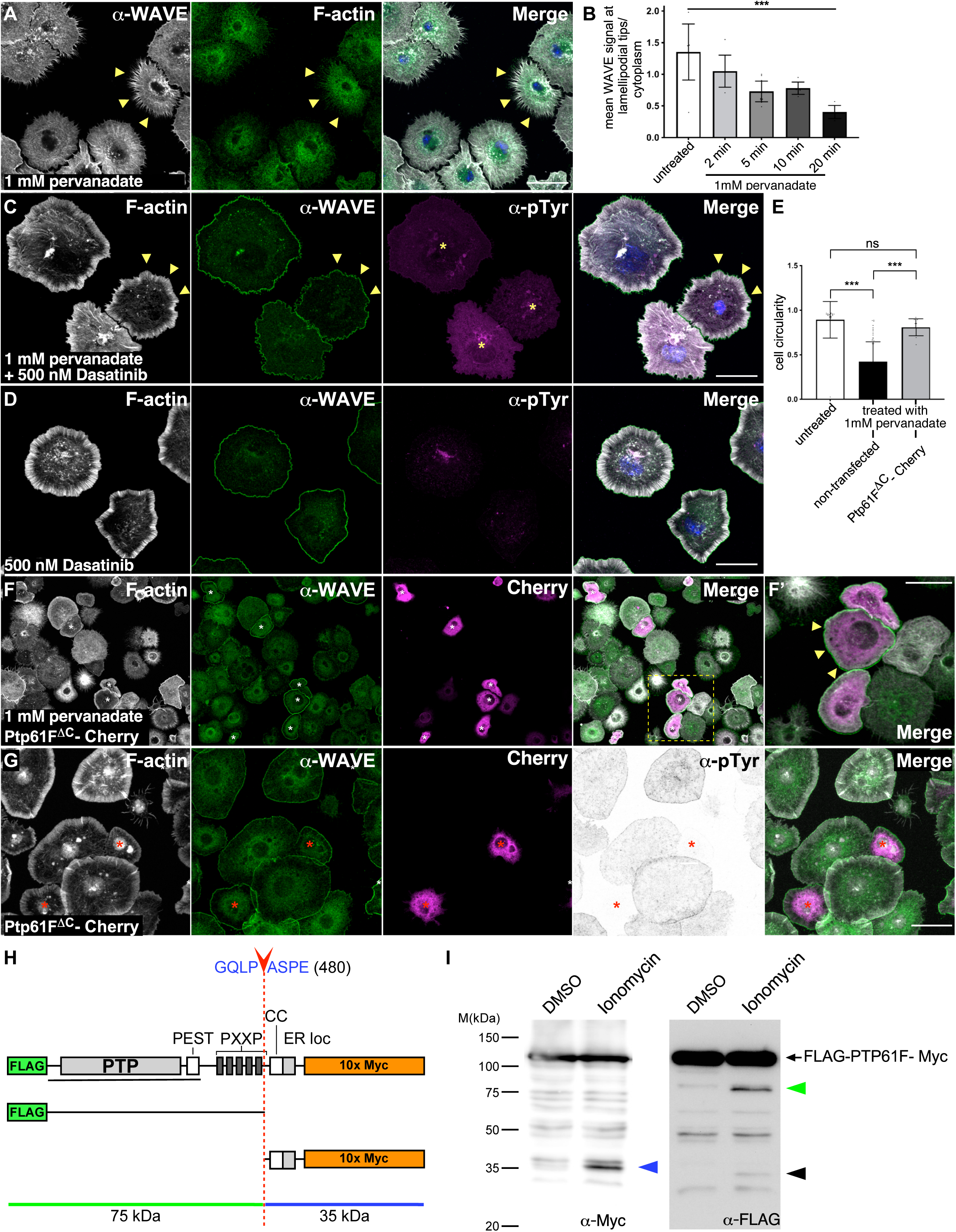
Cytoplasmic PTP61F-ΔC protects cells from pervanadate-induced lamellipodia disruption. **(A)** Treatment of cells with 1 mM pervanadate leads to altered morphology in S2R+ cells, accompanied by a massive reorganization of the actin cytoskeleton and a re-localization of endogenous WAVE from the leading edge to the cytoplasm. **(B)** Quantification of the mean WAVE signal at lamellipodial tips following 1 mM pervanadate treatment over time (in minutes) compared to untreated control cells. Results are the average of three independent experiments. Asterisks indicate significant P-values (*** < 0.001). Error bars represent standard deviation (SD). **(C)** Pervanadate-induced F-actin re-organization and re-localization of endogenous WAVE can be substantially suppressed by co-treatment of cells with 500 nM dasatinib. Note that cells co-treated with pervanadate and dasatinib show a decrease in phosphotyrosine immunostaining (yellow asterisks) different from cells only treated with 500 nM dasatinib **(D)**. **(E, F)** Pervanadate-induced lamellipodia disruption and WAVE re-localization was completely suppressed in cells expressing the cytoplasmic PTP61F-ΔC variant marked by yellow asterisks in F. Results are the average of three independent experiments. Asterisks indicate significant P-values (*** < 0.001). **(G)** Cells expressing PTP61F-ΔC show a striking decrease in phosphotyrosine immunostaining (red asterisks) suggesting that PTP61Fm-ΔC protects cells from pervanadate-induced hyperphosphorylation. **(H)** Scheme of a dual-tagged wild type PTP61F construct containing Flag-tag at the N-terminus and a 10x Myc epitope at the C-terminus. Ptp61F contains a PTP catalytic domain, a PEST motif, five proline-rich motifs (PXXP), a coiled coil and a hydrophobic stretch that is targeted to the ER that localizes PTP to the ER (ER loc). An in silico predicted calpain cleavage site at position 480 (DeepCalpain tool) is marked by a red arrowhead. **(I)** Western blot analysis from lysates of cells transfected with the dual-tagged wild type PTP61F permeabilized with ionomycin to increase intracellular calcium concentrations that activate calpains. N- and C-terminal digestion products of dual-tagged wild type PTP61F were analyzed by using an anti-Myc and an anti-Flag antibody. Flag-PTP61F-Myc has a calculated MW of ∼75 kDa, however, under denaturing conditions in SDS-PAGE the protein migrates above 100 kDa. Accordingly, the corresponding gel electrophoretic mobility deviates for the predicted cleavage N-terminal Fragment (green arrowhead, apparent MW ∼75 kDa versus calculated MW 60 kDa). In addition, a smaller N-terminal fragment is found at 30 kDa suggesting an additional calpain cleavage site (black arrowhead). The C-terminal Myc-tagged fragment including the hydrophobic ER localization sequence appeared at expected 35 kDa (blue arrowhead).

### Calpain mediates the cleavage of PTP61Fm into a C-terminally truncated PTP61Fm-ΔC protein

Like *Drosophila* PTP61Fm, the mammalian ortholog PTP-1B localized to the cytosolic face of the endoplasmic reticulum via its hydrophobic C-terminal targeting sequence (McLaughlin and Dixon, 1993; Woodford-Thomas et al., 1992). Proteolytic cleavage of the hydrophobic C-terminal tail of PTP1B by the calcium-dependent calpain protease is a regulatory mechanism that converts the enzyme from an endoplasmic reticulum (ER)-bound form into a soluble, active cytoplasmic form in human platelets (Frangioni et al., 1993). Once released into the cytoplasm, the C-terminally truncated PTP1B-ΔC has a two-fold higher enzymatic activity (Frangioni et al., 1993) and may have better access to various tyrosine-phosphorylated substrates such as WAVE, thereby regulating its cellular functions in lamellipodia formation. To validate this hypothesis, we analyzed the proteolytic cleavage of the hydrophobic C-terminal tail of PTP61F in S2R+ cells. *Drosophila* has two enzymatically competent calpains, calpain A and calpain B, which contain typical calpain domain structures similar to those of calpains in mammals (Friedrich et al., 2004), and whose activity can be efficiently activated in S2 cell culture by elevated intracellular calcium levels after ionomycin treatment (Bozoky et al., 2009). To analyze a calcium-induced calpain-mediated cleavage of PTP61Fm we generated a PTP61Fm construct with an N-terminal FLAG-tag and a C-terminal Myc-tag (Figure 4). After transfection of the dually tagged PTP61Fm, we used the calcium ionophore ionomycin to induce calcium-dependent cleavage of PTP61Fm. Computational tools such as *DeepCalpain* (http://deepcalpain.cancerbio.info/) predicted several potential calpain cleavage sites in PTP61Fm, including a C-terminal tail domain peptide (aa476-484) containing a single predicted cut site at position 480, in close proximity to the hydrophobic ER localization signal (Figure 3H). Western blotting with antibodies against the N-terminal FLAG tag and the C-terminal Myc tag showed the calcium-dependent appearance of two cleavage products with a molecular weight of 34 and 75 kDa, which corresponds to approximate PTP61Fm fragments cleaved by calpain at position 480 (Figure 3I). Thus, PTP61Fm could be a substrate of calpains which might regulate WAVE-dependent lamellipodia formation.

**Figure 4.**
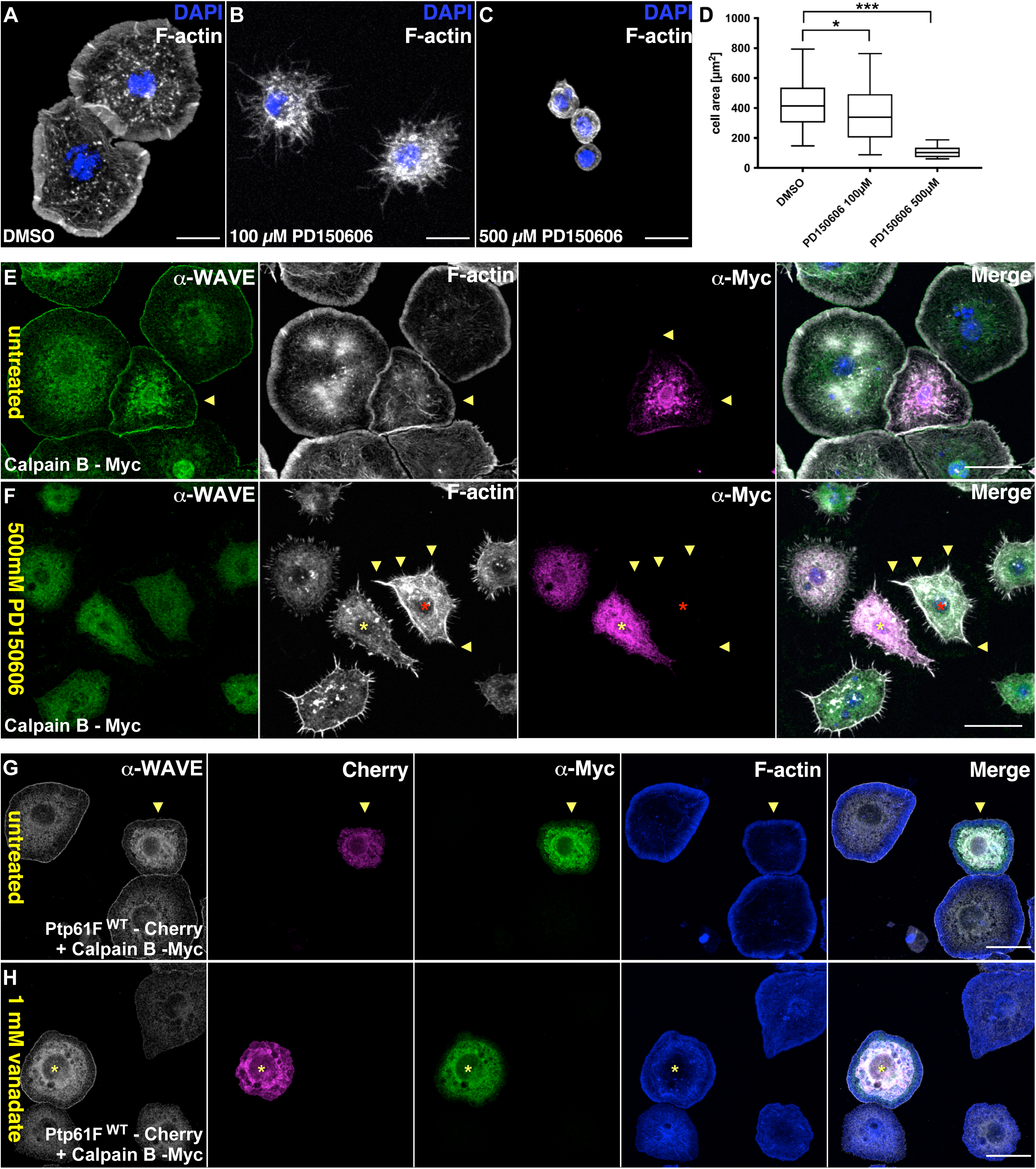
Inhibition of calpain disrupts cell spreading and lamellipodia formation. **(A-C)** Confocal images of S2R+ stained for F-actin (Phalloidin, white) and nuclei (DAPI, blue). Scale bars = 20 µm. **(A)** Untreated cells (DMSO control) spread on ConA with broad circumferential lamellipodia, whereas cells pretreated with **(B)** 100 µM and **(C)** 500 µM calpain inhibitor PD150606 fail to spread and assemble actin-rich lamellipodia. **(D)** Quantification of cell spreading by measuring spread cell area confirmed impaired cell spreading of cells pretreated with PD150606. Cells are significantly smaller than normal and have abnormal Factin networks that completely lack lamellipodia. Results are the average of three independent experiments. Asterisks indicate significant P-values (*< 0.05; *** < 0.001). Error bars represent standard deviation (SD). **(E)** Confocal image of untreated S2R+ cells transfected with a Myc-tagged calpain B construct stained for endogenous WAVE (green), anti-Myc (magenta), F-actin (Phalloidin, white) and nuclei (DAPI, blue). Scale bars = 20 µm. Note that Myc-tagged calpain B colocalized with endogenous WAVE around the nucleus (arrowheads). **(F)** S2R+ cells transfected with a Myc-tagged calpain B were treated with 500 µM PD150606 after complete cell spreading. Lamellipodial protrusion are severely impaired in non-transfected as well as in Myc-tagged calpain B transfected cells (magenta, marked by an asterisk). Endogenous WAVE localization is lost from the leading edge and long, filopodial-like extensions are formed (marked by arrowheads). Scale bars = 20 µm. **(G, H)** Co-expression of Myc-tagged calpain B with a Cherry-tagged-PTP61F-WT construct protects cells from the effects caused by 1 mM pervanadate treatment. Scale bars = 20 µm.

### Inhibition of calpain results in defective WAVE-dependent lamellipodia formation

Genetic inhibition of calpains in NIH-3T3 fibroblasts, specifically by overexpressing Calpastatin results in defective lamellipodia formation and abnormal filopodia induction, indicating that calpain might be indeed involved in actin remodeling during cell spreading (Potter et al., 1998). Calpastatin is an endogenous peptide inhibitor of heterodimeric mammalian calpains that reversibly binds to the small regulatory subunit and inhibits the large catalytic subunit of calpains (Hanna et al., 2008). Different from mammals, *Drosophila* calpains only comprise a large subunit and calpastatin has not been identified in *Drosophila* (Friedrich et al., 2004). However, the activity of both *Drosophila* calpains, calpain A and calpain B, can be efficiently inhibited in S2R+ cell culture using PD150606, a non-competitive calpain specific inhibitor (Bozoky et al., 2009). Interestingly, we found that treatment of S2R+ cells with PD150606 completely blocked S2R+ cell spreading (Figure 4A-D). Control S2R+ cells (DMSO) cells plated on ConA radially spread and adopted a “fried egg-like” morphology with smooth-edged circumferential lamellipodia (Figure 4A). By contrast, PD150606-pretreated cells failed to extend lamellipodial protrusions, and instead cells remained rounded (Figure 4B, C, quantification in figure 4D). Treatment of completely spread cells with the calpain inhibitor PD150606 also led to a rapid collapse of lamellipodia, even in cells overexpressing a Myc-tagged calpain (Figure 4E, F). Interestingly, we found that a Myc-tagged calpain B colocalized with endogenous WAVE around the nucleus (Figure 4E). A prominent co-localization with endogenous WAVE was also observed in cells co-overexpressing Myc-tagged calpain B with a Cherry-taged-PTP61F-WT construct (Figure 4E). Notably, these cells were protected from the effects caused by pervanadate, similar to cells that express the C-terminally truncated PTP1B-ΔC protein (Figure 4E). Thus, this suggest that calpain-mediated proteolytic cleavage of ER-bound wild type PTP61F has a similar protective effect against excessive tyrosine phosphorylation.

### Endogenous *Drosophila* WAVE becomes highly phosphorylated on Y234 modulating WAVE-dependent lamellipodia

To identify WAVE phosphotyrosines, endogenous WAVE was purified from pervanadate-stimulated S2R+ cells by immunoprecipitation using an affinity-purified WAVE antibody (Figure 5A). After a tryptic digest of immunoprecipitates, peptides were analyzed by mass spectrometry. LC-MS analysis revealed a large number of interacting proteins which were coimmunoprecipitated with endogenous WAVE from S2R+ cell, including well-known WAVE binding partners such subunits of the Arp2/3 complex and the WRC (Abi, Sra1, HSPC300), Cip4, Abl but also PTP61Fm (data not shown). Analysis of WAVE phosphopeptides confirmed a major phosphorylation at Y234 with high percentage coverage of 71,3 %, and identified three additional, less common phosphorylation sites in endogenous WAVE, including Y247, Y323 and Y337 (Figure 5B, C; Bodenmiller et al., 2008; Bodenmiller et al., 2007; Chang et al., 2008; Hilger et al., 2009; Hu et al., 2011). To test the role of Y234 on WAVE-dependent lamellipodia formation, we used CRISPR/Cas9 to generate *wave* knock-out cells (*wave* KO) by inserting a GFP expression cassette into the first *wave* exon (codon 24), thereby disrupting its open reading frame (Figure 5C, D). As the result, *wave* KO cells only expressed a small non-functional GFP fusion protein consisting of the first 24 N-terminal amino acids of WAVE and GFP (WAVE 1-24 aa-GFP). This fusion protein could be detected using an anti-GFP antibody and an anti-WAVE antibody, raised specifically against the WHD domain of WAVE (Figure 5D). *wave* KO cells exhibited a characteristic null mutant phenotype with a complete loss of lamellipodia (Figure 5E). Mutant cells showed a prominent stellate morphology, characterized by the emergence of filopodia-like protrusions and an overall strongly reduced circularity (Figure 5E, quantification in Figure 5K). We next performed transient transfections of Myc-tagged WAVE constructs and tested them for their ability to restore lamellipodia formation in *wave*-deficient cells spread on ConA (Figure 5F-I). In these conditions, about 70% of mutant cells could be indeed rescued in lamellipodia formation with WT WAVE, although only half of those lamellipodia were fully developed with a distinct localization of the Myc-tagged WAVE at lamellipodia tips (Figure 5H and I). The remaining cells exhibited either an unchanged stellate cell morphology (no rescue, Figure 5F) or a partial rescue with a predominantly cytoplasmic localization of Myc-tagged WAVE (Figure 5G). Therefore, we quantified overall rescue efficiency by measuring cell circularity. As expected, re-expression of WAVE-WT led to a prominent increase in cell circularity (Figure 5K). Remarkably, rescue with the phosphomutant WAVE-Y234F but not with the phosphomimic WAVE-Y234E significantly further increased cell circularity, almost comparable to S2 Cas9 control cells with normal endogenous WAVE protein level (Figure 5K). Interestingly, such an increased rescue efficiency was even observed upon co-expression of WAVE-WT with a wild type PTP61fm, and particularly when combined with the truncated PTP61f-ΔC construct, which protected cells from increased hyperphosphorylation (Figure 5M). Taken together, these data strongly suggest a model in which calpain-mediated proteolytic release of ER-bound PTP61F antagonizes Abl/Src-dependent WAVE phosphorylation and acts as a mechanism promoting re-localization of WAVE to lamellipodial tips.

**Figure 5.**
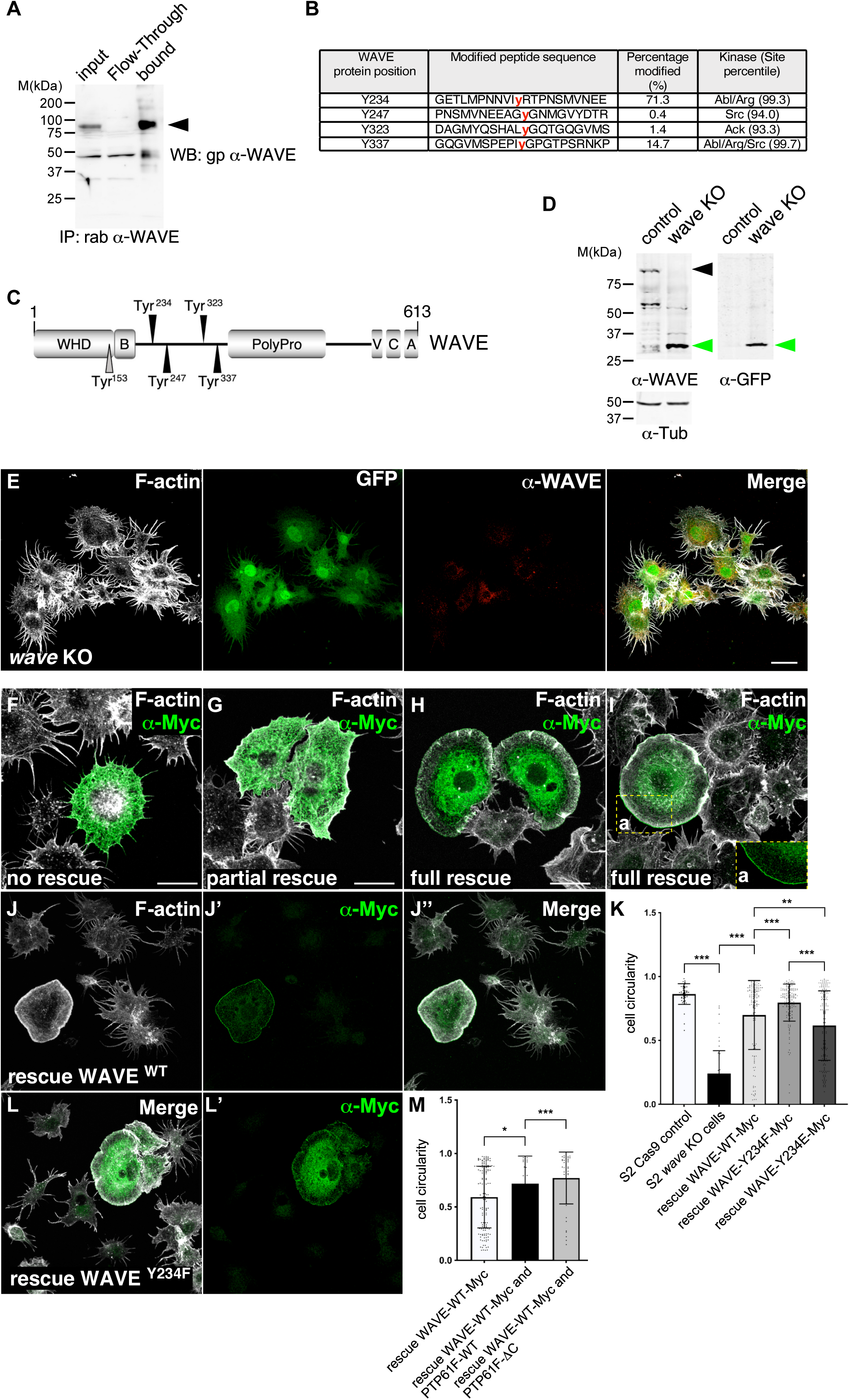
Endogenous *Drosophila* WAVE becomes highly phosphorylated on Y234 modulating WAVE-dependent lamellipodia. **(A)** Immunoprecipitation of endogenous WAVE protein from pervanadate-treated S2R+ cell lysate. **(B)** Mass spectrometry of immunoprecipitated endogenous WAVE protein identifies four phosphotyrosine peptides. The protein position, modified peptide sequence, percentage coverage and predicted kinase are listed. **(C)** Schematic diagram of the domains within WAVE. Phosphotyrosines identified by mass spectrometry are indicated by black arrowheads. The conserved tyrosine residue Tyr153, a major known target for Abl, was not identified (grey arrowhead). **(D)** Western blot analysis from lysates of CRISPR/Cas9 mediated *wave* knock-out cells confirmed a complete loss of endogenous, full-length WAVE protein. Instead, a small non-functional GFP fusion protein consisting of the first 24 N-terminal amino acids of WAVE and GFP (WAVE 1-24 aa-GFP) could by detected by an affinity-purified ant-WAVE-WHD antibody and an anti-GFP antibody at the expected size of 30 kDa. Tubulin served as loading control. **(E)** GFP-expressing *wave* knock-out cells showed a characteristic stellate-cell morphology with a complete loss of lamellipodia. Scale bars = 20 µm. **(F, G, H, I)** Examples of rescue efficiency of *wave* knock-out cells by transient transfection of a Myc-tagged wild type WAVE construct. Note: in **(I)** lamellipodia were fully developed with a distinct localization of the Myc-tagged WAVE at lamellipodia tips. **(J)** *wave* knock-out cell fully rescued by WAVE-WT. **(K)** Quantification of rescue efficiency by measuring cell circularity. Results are the average of three independent experiments. Asterisks indicate significant P-values (*< 0.05; *** < 0.001). Error bars represent standard deviation (SD). S2 Cas9 control n=44, S2 WAVE ko cells n=54, rescue WAVE wt n=142, rescue WAVEY234F n=156 and rescue WAVEY234E n=170. **(L)** *wave* knock-out cells fully rescued by a phosphomutant WAVE-Y234F construct. **(M)** Quantification of increased rescue efficiency (cell circularity) by co-expression of Myc-tagged wild type WAVE with either PTP61F-WT or with PTP61F-ΔC. Results are the average of three independent experiments. Asterisks indicate significant P-values (*< 0.05; *** < 0.001. Error bars represent standard deviation (SD). WAVEmyc plus PTP61fm wt n=26, rescue WAVEmyc plus PTP61fmdC n=44.

## Discussion

Tyrosine phosphorylation-dependent signaling is a critical regulatory mechanism in cells, controlled by the balance between protein tyrosine kinases (PTKs) and protein tyrosine phosphatases (PTPs). Previous studies showed that tyrosine phosphorylation also plays an important role in transmitting extracellular and intracellular signals to the WRC and the actin cytoskeleton (Mendoza, 2013). Tyrosine-dependent WAVE regulation is mainly restricted to the WHD domain, where two prominent tyrosine kinases Src and Abelson (Abl) are reported to phosphorylate WAVE in the meandering region and therefore act on the activity of WAVE. Here, the Abl tyrosine kinase has been described to directly phosphorylate mammalian WAVE2 at the conserved Y150, corresponding to Y151 in WAVE1 and WAVE3 (Krause and Gautreau, 2014; Leng et al., 2005; Sossey-Alaoui et al., 2007; Stuart et al., 2006). In WAVE3, Abl-dependent phosphorylation Y248, Y337 and Y486 need to be additionally modified to completely inhibit phosphorylation (Sossey-Alaoui et al., 2007). Y337 is specific to WAVE3, whereas Y486 is conserved in both WAVE1 and WAVE3, but not in WAVE2 suggesting that phosphorylation of different tyrosine residues in the different WAVE proteins might account for differential roles of the WAVE proteins in mammals (Sossey-Alaoui et al., 2007). Most of these *in vitro* studies suggest that Abl-dependent tyrosine phosphorylation promotes WAVE activation and thus acts positively on Arp2/3-mediated actin polymerization (Krause and Gautreau, 2014; Mendoza, 2013). Src kinases may similarly activate the WAVE complex by phosphorylating Y125 in WAVE1 (Ardern et al., 2006). Structure analysis further supports that phosphorylation Y150/151 but also Y125 interfere with the interaction between the WAVE meander region and CYFIP, releasing the WCA region for Arp2/3 activation (Chen et al., 2010; Ding et al., 2022). However, previous studies on the role of Abl in WAVE-dependent lamellipodia formation have not been investigated further, even though they demonstrate that Abl tyrosine kinase rather inhibits cell migration and restricts the extension of lamellipodia during cell spreading (Jin and Wang, 2007; Kain and Klemke, 2001; Peacock et al., 2007). In this study, we revisited these earlier observations in greater detail. Remarkably, we found that genetic suppression or pharmacological inhibition of Abl and Src did not result in reduced WAVE-dependent lamellipodia formation but rather increases WAVE localization at the leading edge and thus rather promotes lamellipodia formation. By contrast, cells overexpressing a catalytically inactive PTP61F trapping mutant protein phenocopy *wave* mutant cells exhibiting a characteristic stellate-like cell morphology. However, such stellate-like cell morphology could not be observed upon RNAi-mediated depletion of PTP61F suggesting that the trapping mutant protein acts as a dominant-negative mutation also preventing binding of other functional PTPs or adaptors.

The reciprocal regulation of WAVE tyrosine phosphorylation by Abl and PTPs seems to be important for WAVE localization. Tyrosine dephosphorylation of WAVE promotes its localization at the tips of lamellipodia, whereas increased phosphorylation reduces it. This suggests that localization and activity of WAVE might be differentially regulated by different phosphotyrosines. The major phosphorylation sites of the WAVE family (WAVE1-3) differ by specific protein isoforms in mammals, but even in *Drosophila.* Our mass spectrometry analysis confirmed non-conserved phosphorylation site of the *Drosophila* WAVE at Y234 enriched in pervanadate-treated cells as previous reported (Chang et al., 2008). Rescue experiments with a Y234F phosphomutant showed an increased rescue efficiency compared to wild type WAVE protein. Phosphorylation of Y153 in *Drosophila* WAVE, corresponding to Tyr151 in WAVE1/3, Tyr150 in WAVE2 in mammals, could not be detected in our study. However, rescue experiments using a phosphomutant construct show that phosphorylation of Y153 is not essential for WAVE function in lamellipodia formation. On the contrary, phosphomutant Y153F rescued like wild type WAVE with no significant difference (Supplementary figure S1D). Thus, dephosphorylation of WAVE is likely to be an important molecular switch to promote WAVE-dependent lamellipodia formation. Dephosphorylation might an important signal to trigger the re-localization of WAVE from cytosplasmic inactive pools to the leading edge. In *Drosophila* S2R+ cells, a large portion of endogenous WAVE and WRC subunit such as Nap1/Kette is located in the cytoplasm and in perinuclear regions that are thought to be part of the ER (Bogdan and Klämbt, 2003). In plants, the ER can indeed function a reservoir for WAVE/SCAR regulatory complex signaling which is thought to contribute in the early steps of WRC assembly and/or activation (Liu et al., 2021; Zhang et al., 2013). What could drive the release of such mature or premature WRC from such ER reservoir to contribute to WAVE-driven lamellipodia? Our data suggest that calpain-mediated proteolytic cleavage of ER-bound PTP61F might be an important signal that promotes an increase of the pool of active WAVE in lamellipodia. Calpain proteases are known calcium-dependent proteases that cleave numerous substrates in a limited fashion, generating novel activities by substrate proteolysis (Ono et al., 2016; Spinozzi et al., 2021). In *Drosophila*, calpains are restricted to the submembranous compartment close to the ER, a prominent source of calcium for its activation (Friedrich et al., 2004; Vieira et al., 2017). We found that transfected calpain A and B constructs colocalize with perinuclear pools of endogenous WAVE in *Drosophila* S2R+ cells. Ionomycin-induced calpain activation promotes the limited proteolysis of ER-bound PTP61 into a cytoplasmic fragment. Expression of such a truncated PTP-ΔC variant is able to restricts tyrosine phosphorylation of WAVE and protects cells from pervanadate-induced hyperphosphorylation and lamellipodia disruption. Thus, we propose a model in which tyrosine dephosphorylation of WAVE by PTP61F downstream of calpain promotes WAVE-dependent lamellipodia formation. Given that Abl and PTP61F also acts on tyrosine phosphorylation of Kette/NAP1 and Abelson interactor Abi, both central WRC subunits and even a well-known Abl activator and substrate (Chang et al., 2008; Huang et al., 2007; Ku et al., 2009; Stephan et al., 2008), we propose an additional regulatory feedback mechanism by Abi that modulate the stability and localization of WRC.

### Experimental procedures

#### Plasmid construction and mutagenesis

The open reading frame (ORF) of the endoplasmic reticulum (ER)-bound form of PTP61F (PTP61Fm), corresponding to amino acids 1–339, was amplified by PCR from cDNA LP02164 (BDGP) and cloned into the pENTR vector using TOPO cloning (Gateway system, Invitrogen). The resulting entry clones were recombined into pUAST expression vectors (Gateway) containing various N- or C-terminal tags. The substrate-trapping mutant (PTP61fm D203A) was generated by site-directed mutagenesis using primers carrying the desired point mutation, followed by DpnI digestion. Phospho-deficient and phospho-mimetic WAVE mutant constructs were generated using standard PCR-based mutagenesis protocols. The ΔC construct (pENTR) was kindly provided by S. Önel. All ORFs with or without a STOP codon were cloned into pENTR gateway-entry vector (pENTR/ D/ TOPO) and additional tagging (GFP, myc, FLAG) was introduced by Gateway cloning as required. All constructs were verified by sequencing (Seqlab).

#### Cell culture, transient transfection, immunoprecipitation, and immunoblotting

*Drosophila* S2R+ cells (Yanagawa et al., 1998) were maintained in 1× M3-Schneider medium (Gibco) supplemented with 10% fetal bovine serum and Pen/ Strep (herein called M3++) at 25°C.For transient transfection with plasmid, FuGENE HD (E2312, Promega) was used as a vehicle, following the manufacturer’s directions. For RNA interference (RNAi), double-stranded RNA (dsRNA) was added to serum-free medium. After 2 h, complete medium was added and cells were incubated for 4–5 days.

#### dsRNA Preparation

For RNAi-mediated knockdown experiments, double-stranded RNA (dsRNA) was added to serum-free culture medium, according to the established protocol (Bogdan and Klämbt, 2003). Briefly, 1 µg dsRNA was added to the cells in M3—for 1 hours, followed by medium change and incubation for 4-5 days. Target sequences were amplified by PCR using primers containing T7 promoter sequences. dsRNA was synthesized by in vitro transcription using T7 RNA polymerase (HiScribe T7 RNA Synthesis Kit, NEB) according to the manufacturer’s protocol.

#### CRISPR/Cas9-Mediated Genome Editing

S2-Cas9 cells were used to generate a WAVE knockout cell line as previously described (Lenz et al., 2025). Briefly, to promote homologous recombination, cells were pre-treated with dsRNA targeting Lig4 and mus308. Cells were co-transfected with a plasmid encoding sgRNA under a U6 promoter and a donor construct containing a GFP tag and antibiotic resistance cassette flanked by homology arms. Homology-directed repair constructs were amplified from pMH3 (Addgene #52528). Selection was performed using 10 µg/ml blasticidin for at least 14 days. Successful integration was verified by genomic PCR.

#### Protein Extraction and Western Blotting

Cells were lysed in lysis buffer containing 20 mM Tris (pH 7,5), 150mM NaCl, 1mM EDTA, 1mM EGTA, 1% Triton x-100 with protease and phosphatase inhibitors (2,5 mM Natrium-Pyrophosphat, 1 mM β-Glycerolphosphat, mM NaVa3). Lysates were incubated on ice and cleared by centrifugation. Protein concentrations were determined using Bradford assays (Bio-Rad). Equal amounts of protein were separated by SDS-PAGE and transferred to activated PVDF membranes (Thermo scientific). PVDF membranes were blocked in 5% milk in TBS-T and incubated with primary antibodies overnight at 4°C, followed by HRP-conjugated secondary antibodies. Detection was performed using chemiluminescence (ECL).

#### Immunoprecipitation experiments

For immunoprecipitation, lysates were incubated with specific antibodies followed by Protein A/G agarose beads. Alternatively, Myc-tagged proteins were isolated using Myc-Trap (ChromoTek) according to the manufacturer’s instructions. Bound proteins were eluted in Laemmli buffer and analyzed by Western blot.

#### Immunofluorescence staining

Cells were plated on concanavalin A-coated coverslips, fixed with 4% paraformaldehyde, and shortly permeabilized with Triton X-100. Primary antibody incubation was performed, after blocking, for 2h, followed by incubation with fluorophore-conjugated secondary antibodies. F-actin and nuclei were stained with phalloidin and DAPI, respectively. Images were acquired using Leica confocal microscopes (SP8) and processed with ImageJ/FIJI. The following primary antibodies were used: anti-WAVE (affinity-purified guinea pig and rabbit (Bogdan et al., 2005); anti-Myc 9E10 (DSHB hybridoma supernatant); anti-phosphotyrosine pTyr (1:1000; Cell signaling). Secondary antibodies conjugated to Alexa fluorophores were used according to standard protocols.

#### Image analysis

Cell size and fluorescence intensity were quantified using ImageJ/Fiji (NIH). Regions of interest (ROIs) corresponding to whole-cell boundaries were manually defined using the freehand selection tool based on cortical actin staining. Mean fluorescence intensities were measured for both cortical and cytoplasmic compartments. To quantify cortical signal, the cell periphery was manually traced and the corresponding mean fluorescence intensity was determined. For cytoplasmic measurements, the cortical selection was converted into an area ROI and adjusted to exclude the cell cortex, allowing determination of mean cytoplasmic fluorescence intensity. Lamellipodial-to-cytoplasmic intensity ratios were subsequently calculated for each cell. In addition, cell shape parameters, including circularity, were extracted from the defined whole-cell ROIs.

#### Pharmacological treatments

Cells were treated with the following compounds: calpain inhibitor PD150606 (100–500 µM, Sigma-Aldrich); ionomycin (8 µM) + CaCl₂ (10 mM); Dasatinib and Imatinib (various concentrations as indicated). Calpain inhibitor PD150606 (Sigma-Aldrich, Merck) was diluted to 100µM or 500µM in M3++ and either added during or after cell spreading on ConA for 30 minutes. For calpain activation cells were incubated with 8 μM Ionomycin (Sigma Aldrich) and Cacl2 (10mM) for 30 minutes before cell lysis. Pervanadate was freshly prepared by mixing sodium orthovanadate with H₂O₂ and applied to cells in M3 withous FCS for 20–30 min.

#### Protein Digestion and Sample Preparation for mass spectrometry

Proteins were eluted from the sepharose beads by the addition of 30 ul of 2% SDS and incubation at 90°C for 10 minutes with shaking. Afterwards the beads were shortly pelleted by centrifugation and supernatant was transferred to a fresh, 500 ul Eppendorf tube. The elution procedure was repeated for the second time and both supernatants were combined. Next, proteins were subjected to mass spectrometry–based proteomic analysis using a modified single-pot solid-phase–enhanced sample preparation (SP3) protocol. This workflow integrates protein solubilization, enzymatic digestion, detergent removal, and peptide cleanup into a single, streamlined procedure prior to LC–MS analysis. Protein capture was performed by binding samples to paramagnetic beads in a solution of 70% (v/v) anhydrous acetonitrile (ACN) under neutral pH conditions. To eliminate impurities, the beads underwent sequential washes using 70% ethanol and 100% anhydrous ACN. Following the near total evaporation of ACN, the beads were resuspended in 30 µL of 50 mM TEAB. Proteolysis was initiated by adding 1 µg of trypsin (Promega), with the reaction proceeding overnight at 37 °C under constant agitation. Post-digestion, a SpeedVac concentrator was used to reduce sample volumes to roughly 5 µL. Peptides were then re-immobilized on the beads by increasing the ACN concentration to >95% (v/v), followed by two additional washes with the same solvent. The peptides were eluted in 50 µL of 0.1% formic acid supplemented with 0.015% (w/v) n-dodecyl-β-D-maltoside (DDM) and transferred to MS-certified vials. Peptide concentrations were quantified via the Pierce Fluorometric Quantitative Peptide Assay (Thermo Scientific), and sample concentrations were normalized to 25 ng/ul to ensure consistent loading for mass spectrometry.

#### LC–MS/MS Configuration

Peptide analysis was conducted on a timsTOF Ultra mass spectrometer integrated with a nanoElute nanoLC system (Bruker Daltonics). For each run, approximately 50 ng of peptide material was initially trapped and separated on a 25 cm PepSep Ultra C18 analytical column (1.5 µm particle size, 75 µm ID). Separation was achieved at a flow rate of 300 nL/min using a graduated mobile phase gradient (Solvent A: 0.15% formic acid in water; Solvent B: 99.85% ACN, 0.15% formic acid). Peptides were eluted using following gradient: 2% to 35% solvent B over 90 min. The analytical column was linked to the mass spectrometer through a 20 µm CaptiveSpray emitter. Data acquisition utilized a data-dependent acquisition (DDA) strategy with a default dda method provided by Bruker. In short: full MS1 scans were captured at 45,000 resolution (m/z 100–1,700). For ion mobility, a 1/K₀ range of 0.70–1.45 V·s/cm² was employed with a 100 ms ramp time, with a cycle time of 0.53 s.

#### MS Data Processing and Database Search

MS data were processed and analyzed using MaxQuant (v. 2.5.1.0). Raw files were searched against a *Drosophila* UniProt database (38,869 entries; October 2024). The search was performed using the TIMS-DDA configuration, with Trypsin/P specified as the proteolytic enzyme allowing for a maximum of two missed cleavages. Carbamidomethylation of cysteine was set as a fixed modification, while methionine oxidation, protein N-terminal acetylation, and phosphorylation (STY) were included as variable modifications. Precursor mass tolerances were set to 20 ppm for the initial search and 10 ppm for the main search, while the fragment mass tolerance for the TOF detector was maintained at 25 ppm. Peptide identifications were restricted to a minimum length of 7 amino acids. A false discovery rate (FDR) of 1% was applied at the PSM, protein, and site levels to ensure high confidence in the results. For quantification, the Match Between Runs (MBR) feature was enabled with a matching time window of 0.4 minutes and an ion mobility window of 0.05 1/K₀. Protein groups were identified requiring at least one razor or unique peptide.

## Acknowledgements

We thank the Bloomington Stock Center and VDRC for fly stocks. We thank Peter Graumann and Witold Szymanski for mass spectrometry analysis (Core Facility for Mass Spectrometry & Proteomics, University Marburg). We thank Oliver Hantschel for sharing imatinib. We thank Doris Wagner for S2R+ cell culture experiments and Susanne Önel for providing a PTP61Fm-ΔC construct cloned into pENTR. We thank Dennis Klug, Katja Rust and Mila Y. Höhne for critical reading of the manuscript. The work was supported by grants to S.B. (BO1890/5-1 and BO1890/7-1) from the Deutsche Forschungsgemeinschaft (DFG).

## Author contributions

S.B. designed the project. S.B. made the figures and wrote the manuscript. W.M. performed the experiments and the statistical analysis A.H. cloned the PTP61fm constructs.

---

**Supplementary figure S1.**
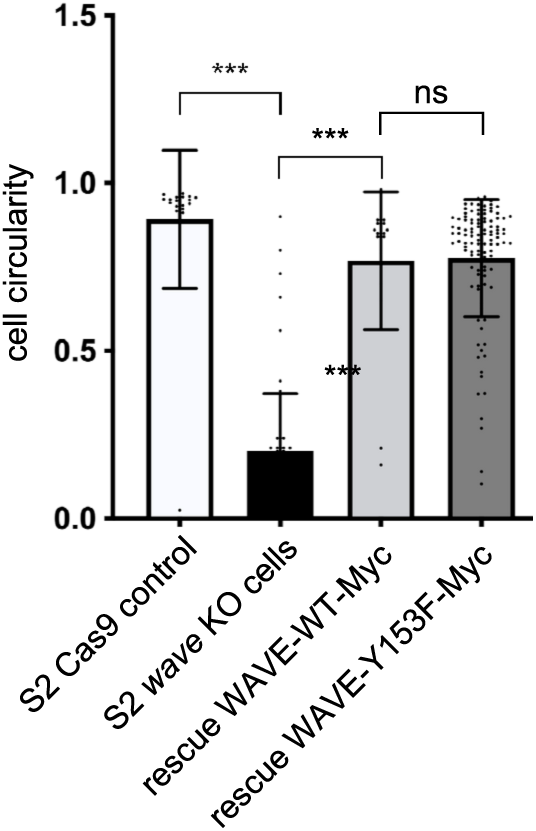
**(A, B)** RNAi-meditated depletion of *PTP61F* affects neither in S2R+ cell morphology nor WAVE localization with no significant differences compared to cells treated with double stranded RNA directed against a luciferase control (*luc* RNAi). **(C)** RNAi specificity was confirmed by using a transfected PTP61F-mCherry fusion which was efficiently downregulated in contrast to control proteins like Tubulin. **(D)** Phosphorylation of Y153 is not essential for WAVE function in lamellipodia formation Quantification of rescue efficiency of *wave* KO cells re-expressing a phosphomutant Y153F construct by measuring cell circularity. Circularity is given as 4p(cell area/cell perimeter^2^) and describes the similarity of an object to a perfect circle. Results are the average of three independent experiments. Asterisks indicate significant P-values (*** < 0.001; ns: no significant difference). Error bars represent standard deviation (SD). S2 Cas9 control n=44, S2 WAVE ko cells n=54, rescue WAVE wt n=142, rescue WAVEY153F n=109.

